# Cytokines and schizophrenia revisited: a two-sample multi-marker Mendelian randomization approach

**DOI:** 10.1101/870022

**Authors:** Hongyan Ren, Yajing Meng, Yamin Zhang, Qiang Wang, Wei Deng, Xiaohong Ma, Liansheng Zhao, Xiaojing Li, Yingcheng Wang, Pak Sham, Tao Li

## Abstract

**Background:** Schizophrenia is a complex mental disorder with recent evidence suggesting a critical immune component underpinning its pathophysiology. Two-sample Mendelian randomization (MR) provided an opportunity to probe the immune changes in schizophrenia by harnessing the increasing availability of summary-level data from large GWAS consortia.

**Objective:** To map the extensive immune response of schizophrenia in terms of cytokines/chemokines and to explore the effect of cytokines induced by schizophrenia (SCZ-induced cytokines) on the brain structure and function

**Sources and methods:** Using the summary-level data generated from GWAS of schizophrenia, cytokines in the peripheral blood and imaging-derived phenotypes (IDPs), we performed two rounds of two-sample MR analysis; the identified cytokines from first round of analysis (schizophrenia => cytokines) were modeled for its underlying structure and subsequent clustering analysis further grouped SCZ-induced cytokines based on their genetic similarities. The multi-phenotype summary statistics of each cytokine module were then used as instrumental variables (IVs) for the second round of MR analysis to detect their effect on brain structure and function.

**Results:** The first round of MR analysis identified nine cytokines, the highlight of which includes IL18 (OR = 1.292, P = 8.37 × 10^−42^) and TNFa (OR = 0.721, P = 7.33 × 10^−6^), to be causally associated with schizophrenia. These SCZ-induced cytokines could be clustered into three modules. The second round of MR analysis (cytokine module => IDPs) indicated that module B (SCGFb-IP10-CTACK-IL6) significantly increased the level of IDPs including IDP_T1_SIENAX_peripheral_grey_normalised_volume (β = 0.0453, P = 4.40×10^10^), IDP_dMRI_TBSS_MD_Posterior_corona_radiata_R (β= 0.0584, P = 8.89× 10^−16^) and IDP_dMRI_TBSS_MD_Cingulum_hippocampus_R (β = 0.0563, P = 9.88× 10^−15^), with module C (IL18-GROa-TNFa) increasing the level of IDP_dMRI_TBSS_L2_Posterior_thalamic_radiation_R (β= 0.0341, P = 2.67× 10^−6^).

**Conclusion:** Our study, for the first time, mapped the causal link from schizophrenia to the comprehensive immune responses, and the findings suggest immune networks play a role in pathophysiology of schizophrenia by mediating the deviations of total gray matter volume and white matter fibers possibly in the mesolimbic system.

## Introduction

Schizophrenia is a complex disorder with a genetic architecture involving both polygenicity and pleiotropy (Consortium, 2013; Purcell et al., 2014). Before the advance and widespread application of genetic decoding at the genome level, almost all the hypothesis related to the pathophysiology of schizophrenia came from the post-hoc validation following the serendipitous discovery of treatment agents, which lacks precision with a mixed response (Delay, Deniker, & Harl, 1952; Gründer & Cumming, 2016). With technological breakthroughs and cost reduction of genome-wide genotyping, the psychiatric consortia, such as psychiatric genetic consortium and iPSYCH, conducted genome-wide association studies (GWASs) of schizophrenia in a large population scale (Pardiñas et al., 2018). The findings from these studies cast some novel lights on the biological underpinnings of schizophrenia, the most notable and replicated of those lights should be the implication of immune dysfunction in the pathogenesis of schizophrenia. Although relevant hints have been provided in the past by the case report, anecdote, and post-mortem studies, the recently published GWASs, for the first time, confirmed such involvement based on the large population sample (Van Kesteren et al., 2017). However, the precise mapping of the immune repercussions of schizophrenia remains elusive. Many studies tried to capture the change in immune response to schizophrenia, for example, change of cytokines in peripheral blood of patients with schizophrenia, with inconsistent findings. The reasons behind such an inconsistency can be two-fold, firstly, schizophrenia is highly heterogeneous in clinical manifestations, converging lines of evidence have indicated that different cluster of symptoms, e.g. positive symptoms, negative symptoms or cognitive impairment, might be underpinned by distinct psychopathological process; secondly, the majority of studies on immune dysfunction of schizophrenia focused on one or two components of immune function, for example, the change of one or a few cytokines. Under the assumption of omnigenic model, in the cell types that are relevant to a disease, it appears that essentially all genes contribute to the condition. In the context of such, schizophrenia should induce immune change in a network manner involving multiple components, such as cytokines.

While it is time- and resource- consuming to collect relevant information on the immune changes in patients with schizophrenia in a large scale of population, two-sample Mendelian randomization (MR) which leverages the publicly accessible summery-level data from the genome-wide association studies (GWAS) provides a valuable avenue to detect the causal relationship between phenotype pairs, and gains popularity due to the immutability of genetic variants as instrumental variables (IVs) and the free accessibility of a vast resource of GWAS results of different phenotypes and traits. Unlike one-sample MR, two-sample MR does not need the effect of the IV-risk factor association and the IV-outcome factor association to be collected in the same sample of participants, which could evade the confoundings such as “winner’s curse”; further, unlike one-sample MR, the weak IVs in the two-sample MR bias the result to the null (Lawlor, 2016). Hartwig *et al.* adopted such an approach to draw a quantitative causal link from CRP (C-reactive protein) and IL1 (interleukin 1) to the lifetime risk of schizophrenia (Hartwig, Borges, Horta, Bowden, & Smith, 2017). Although the study is rigorous with a notable impact, there are a few limitations the authors have yet to address; first, Hartwig *et al.* used the genetic variants passing the genome-wide significance threshold (5×10^−8^) of association as IVs, these IVs could only explain a small proportion of the phenotypic variance of complex traits. Studies using polygenic risk score (PRS) have shown that including more SNPs associated at a subthreshold level could increase the predictive power; second, the study only focused on a few candidate cytokines. Reviews by Kroken *et al.* indicated that there might be one or a few cytokine networks underlying the syndromic symptoms of schizophrenia (Kroken, Sommer, Steen, Dieset, & Johnsen, 2019).

Our study seeks to use two rounds of two-sample MR analysis (schematics of study design shown in Figure 1) to answer the following two questions: (1) What is the immune response to schizophrenia in terms of cytokines/chemokines? (2) What is the effect of those cytokines responsive to schizophrenia on the brain structure and function?

**Figure 1.**
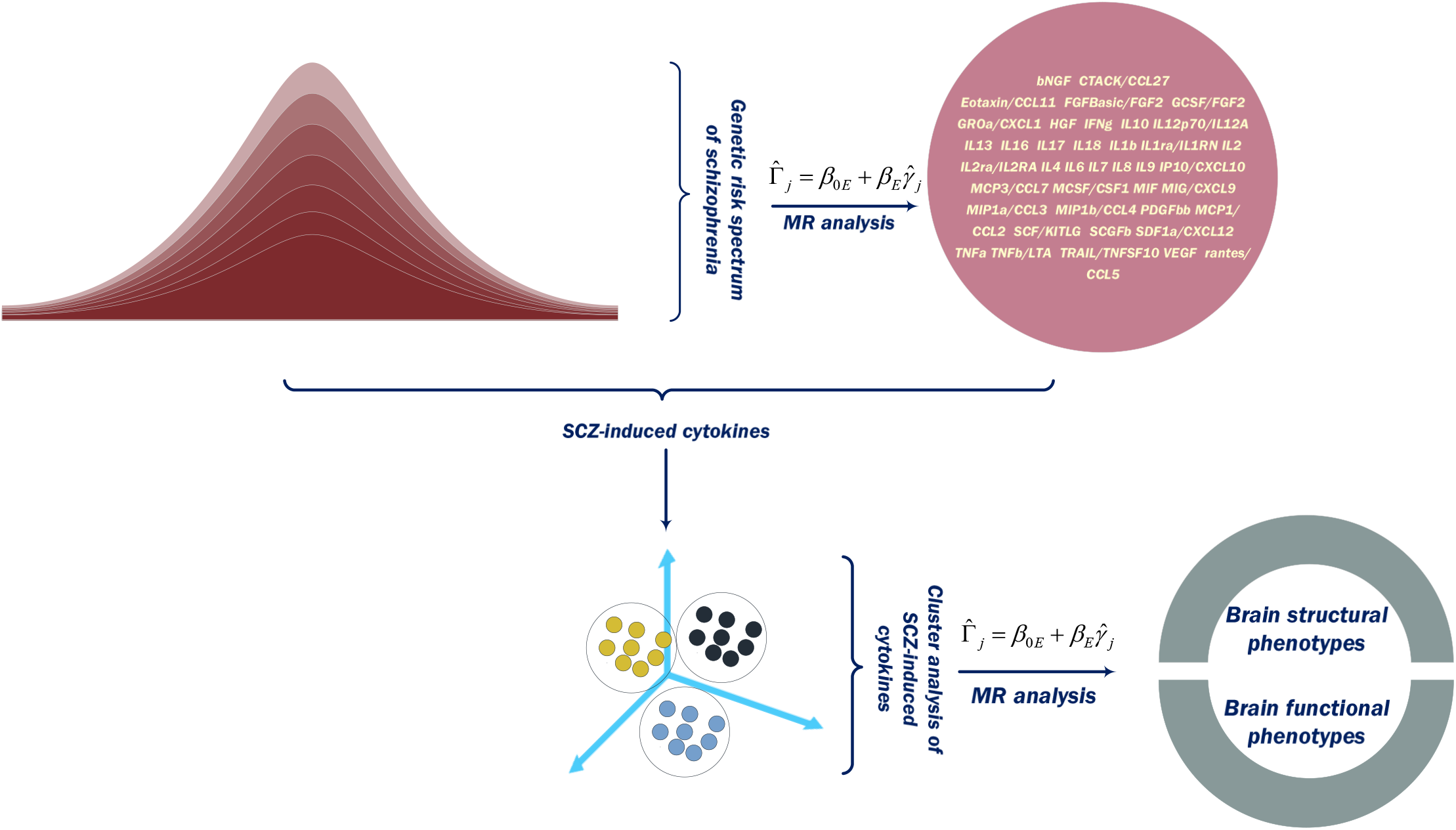
Schematic diagram of our data analysis strategy for two rounds of Mendelian randomnization studies

## Methods

### Data source

For the first-round of MR analysis, our study used the summary statistics of schizophrenia GWAS conducted by Pardina et al. in 11,260 cases and 24,542 controls of European ancestry (Pardiñas et al., 2018) as the exposure. For the immune outcomes, we used the summary statistics of GWAS of 40 cytokines/chemokines in the peripheral blood of 8500 healthy Finnish volunteers (Ahola-Olli et al., 2017). No sample overlapping was identified between two datasets.

### Instrumental variable (IV) selection

To capture the immune changes of schizophrenia in a gradient manner, we chose the SNPs associated with schizophrenia at seven different P-value thresholds (5× 10^−8^, 1 × 10^−7^, 1 × 10^−6^, 1 × 10^−5^, 1 × 10^−4^, 1 × 10^−3^, 1 × 10^−2^) as the instrumental variables (IVs). Before the MR analysis, all IVs were clumped using the following parameters (--clump-p1 1 --clump-p2 1 --clump-r2 0.001) to ensure independence (LD equilibrium) between them.

### MR analysis

Many methods have been developed to conduct MR analysis, each tackling specific issues of MR. However, only recently have methods been tailored to the challenges facing two-sample MR in the context of complex traits, such as measurement error due to weak IVs, widespread horizontal pleiotropy in the genome, violation of InSIDE (INstrument Strength Independent of Direct Effect) assumption. Given the design of our current study, we chose two recently developed algorithms, MR-RAPS and MRMix, which alleviate the above-mentioned issues and provide less-biased causal estimates when including multiple markers as IVs. In brief, MR-RAPS assigns weights to IVs according to their strength by using a Bayesian approach with informative prior for the weighting scheme (Zhao, Wang, Hemani, Bowden, & Small, 2018), and is robust to deviations of usual IV assumptions. MRMix took advantage of a working parametric model for the underlying bivariate effect size distribution of the SNPs across pairs of traits, which allows genetic correlation to arise both from causal and noncausal relationships; the extensive simulation studies showed that MRMix could provide much better trade-off between bias and variance than existing estimators in a broad set of scenarios (Qi & Chatterjee, 2019). In the sensitivity analysis, we employed MR-Egger and weighted mean to analyze the same harmonized data. The analysis was conducted using packages “TwoSampleMR”, “mr.raps” and “MRMix”in the R. We chose cytokine with a causal estimate surviving Bonferroni correction for multiple comparisons (0.05/43 = 0.0012) at any of seven P-value thresholds for IV selection as the SCZ-induced cytokine.

Since the exposure studied here is a binary variable, the GWAS of which was carried out in the context of the log-odds scale. For the sake of interpreting the results, we converted the causal estimates by multiplying 0.693 (ln 2), followed by exponentiation to represent the change in outcome (cytokines) as per 2-fold change in the polygenic burden of schizophrenia.

### The detection of the potential structure underlying SCZ-induced cytokines

The above step identified nine cytokines to be responsive to schizophrenia. To explore the inner structure among nine SCZ-induced cytokines, we implemented a genomic SEM to detect the potential commonality underlying the nine cytokines. If such a commonality failed to be identified, we took advantage of the genetic correlation matrix of these nine cytokines, generated from LD score regression, for the agglomerative hierarchy clustering analysis (Zhang, Murtagh, Van Poucke, Lin, & Lan, 2017). The analysis first computed the proximity matrix, in our study we used the Manhattan distance to create the matrix; let each data point as a cluster at the beginning, then we merged the two closest clusters and update the proximity matrix until only a single cluster remains. To test the robustness of preliminary clustering results, we bootstrapped the clustering results for 10000 times to assess the uncertainty of the clustering results, quantified by the P values indicative of how strong the cluster is supported by data. The analysis was conducted in the R.

### Potential impact of SCZ-induced cytokines on brain imaging phenotypes

To reveal the phenotypic consequences of SCZ-induced cytokine modules, especially their effect on the brain structure and function, we used the results of cluster analysis from the previous step to conduct the second round of exploratory two-sample MR analysis with the IDP (imaging-derived phenotype) from the UK biobank as the outcomes (Elliott et al., 2018). We included 627 IDPs covering different imaging modalities in our current MR analysis. For more details of IDPs used here, refer to https://www.fmrib.ox.ac.uk/ukbiobank/gwas_resources/index.html. Prior to the analysis, we implemented a multi-trait analysis using MTAG to generate new composite summary statistics for each cytokine module. The SNPs associated with cytokine cluster at a lenient genome-wide significance level of 0.00005 were chosen as IVs, and the analysis was carried out using MRMix.

## Results

### Multiple cytokines were implicated in the immune response induced by schizophrenia, with a modest effect size

Prior to carrying out the MR analysis, we calculated the genetic correlations (*rg*) for 40 cytokines and schizophrenia using the LD score regression. The only cytokine correlated with schizophrenia at the nominal significant level was IFN*g* (*rg* = 0.3204, P = 0.02298). Besides, some trends of correlation were found between TNFa (*rg* = 0.01783), IL2 (*rg* = 0.5882), IL6 (*rg* = −0.2277), IP10/CXCL10 (*rg* = −0.5287), IL8 (*rg* = 0.2994), rantes/CCL5 (*rg* = 0.3166), and schizophrenia, although none of them could reach the level of nominal significance (Figure 2).

**Figure 2.**
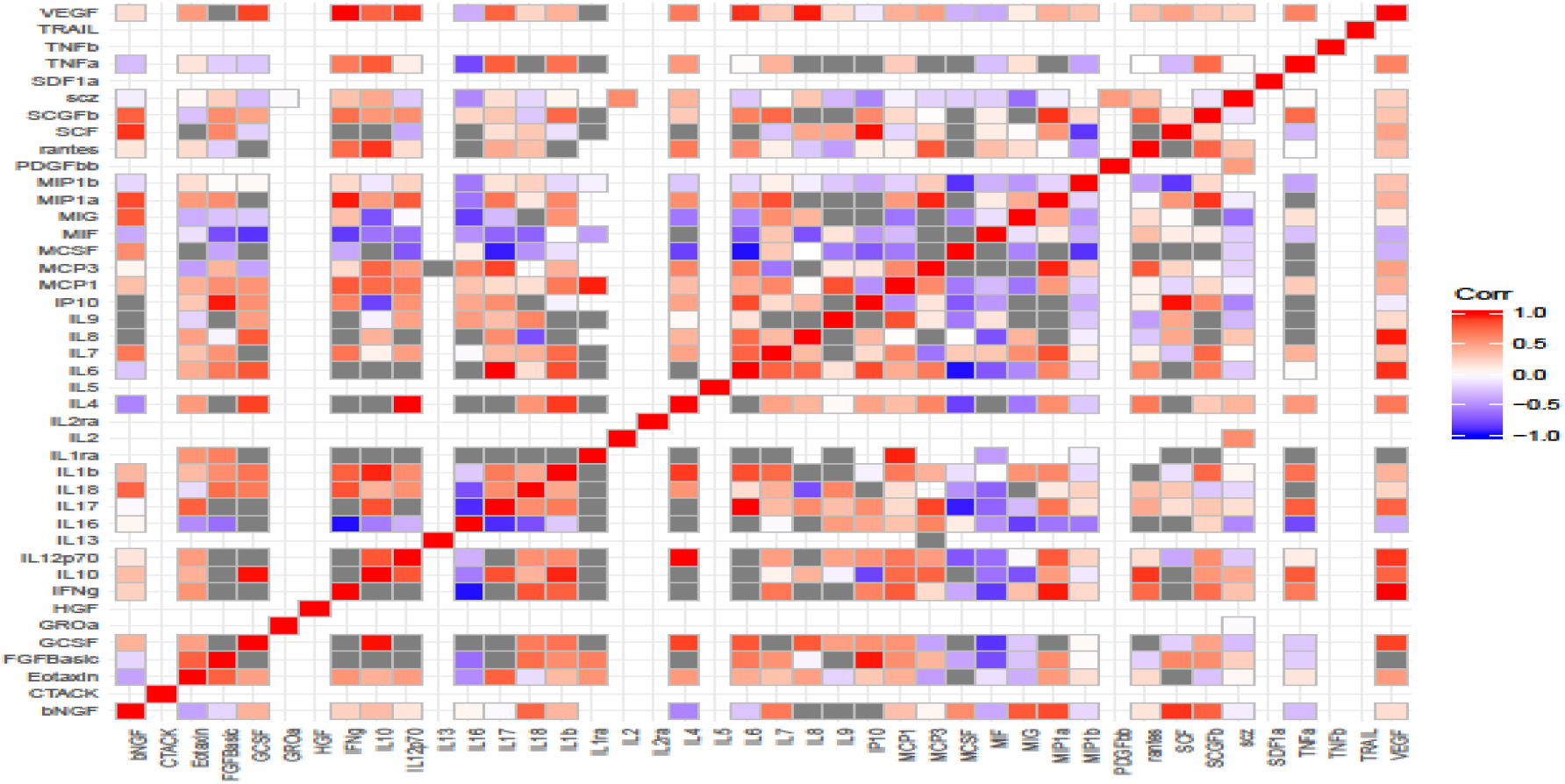
Genetic correlation between 43 cytokines and between each cytokines and schizophrenia using LDSC regression

The results of MR analysis were shown in Table 1; nine cytokines were identified to be SCZ-induced cytokines. The pattern of causal estimates for different cytokines implies their different underlying genetic components.

**Table 1.** Estimates for causal effects (one unit change of cytokine as per 2 SD-unit increase in risk burden of schizophrenia) of various putative risk-factors on cytokine outcomes

|  |  | P value threshold<br>(N) |  |  |  |  |  |  |
| --- | --- | --- | --- | --- | --- | --- | --- | --- |
|  |  | 0.00000005 | 0.0000001 | 0.000001 | 0.00001 | 0.0001 | 0.001 | 0.01 |
|  |  | 121 | 133 | 224 | 363 | 463 | 682 | 777 |
| GROa | MR-RAPS<br>(shrinkage) | 0.9031 | 0.9107 | 0.877* | 0.895* | 0.925** | 0.950* | 0.9553 |
|  | MR-Egger | 1.0728 | 0.9126 | 1.002 | 0.9443 | 0.9596 | 0.9818 | 0.9885 |
|  | Weighted median | 0.8645 | 0.8827 | 0.891 | 0.9151 | 0.9045 | 0.907 | 0.913 |
|  | MRMix | 0.717 | 0.727 | 0.712 | 0.735 | 0.796 | 0.795 | 1.256*** |
| IL6 | MR-RAPS<br>(shrinkage) | 0.9606 | 0.9669 | 0.94 | 0.934* | 0.9917 | 0.9887 | 0.9907 |
|  | MR-Egger | 1.0461 | 1.0287 | 1.1123 | 0.9653 | 1.0353 | 1.0294 | 1.0307 |
|  | Weighted median | 0.9898 | 0.9921 | 0.9657 | 0.9577 | 0.9757 | 0.9757 | 0.9759 |
|  | MRMix | 1.172 | 1.283 | 1.042 | 1.101* | 1.247* | 1.414 | 1.385** |
| IL9 | MR-RAPS<br>(shrinkage) | 0.874* | 0.8989 | 0.876* | 0.885** | 0.941* | 0.9724 | 0.9853 |
|  | MR-Egger | 1.2826 | 1.0325 | 1.073 | 0.9793 | 0.9224 | 0.9756 | 0.9485 |
|  | Weighted median | 0.969 | 1.0186 | 0.9723 | 0.971 | 1.0095 | 1.024 | 1.0225 |
|  | MRMix | 1.324 | 1.296 | 0.986 | 0.8832* | 1.268 | 1.256 | 0.972 |
| IP10 | MR-RAPS<br>(shrinkage) | 0.9366 | 0.9542 | 0.9343 | 0.907* | 0.909**** | 0.935*** | 0.954* |
|  | MR-Egger | 0.6908 | 0.7323 | 0.888 | 1.0189 | 0.9859 | 0.8415 | 0.8647 |
|  | Weighted median | 0.9901 | 1.0074 | 0.98 | 0.9353 | 0.93 | 0.9325 | 0.9461 |
|  | MRMix | 1.003 | 1.003 | 0.907 | 1.021 | 1.163 | 1.042 | 0.732 |
| CTACK | MR-RAPS<br>(shrinkage) | 0.856* | 0.856* | 0.866** | 0.9123 | 0.9994 | 0.9822 | 0.9912 |
|  | MR-Egger | 0.6592 | 0.6467 | 0.6527 | 0.6631 | 0.8263 | 0.9187 | 0.8472 |
|  | Weighted median | 0.8582 | 0.8586 | 0.8683 | 0.9161 | 0.9385 | 0.9713 | 0.9708 |
|  | MRMix | 1.197 | 1.197 | 1.028 | 0.979 | 0.979 | 1.042 | 1.035 |
| TNFa | MR-RAPS<br>(shrinkage) | 0.879* | 0.893 | 0.8999 | 0.9272 | 1.0443 | 1.0216 | 1.0066 |
|  | MR-Egger | 1.6921 | 1.4671 | 1.3006 | 1.1958 | 1.1231 | 0.997 | 1.0961 |
|  | Weighted median | 0.8818 | 0.8883 | 0.9198 | 0.9442 | 0.9657 | 1.0051 | 1.0115 |
|  | MRMix | 0.724 | 1.218 | 0.712**** | 0.9205 | 0.718 | 1.356 | 1.414 |
| SCGFb | MR-RAPS<br>(shrinkage) | 1.039 | 1.0632 | 1.0122 | 1.0443 | 1.066* | 1.059** | 1.0347 |
|  | MR-Egger | 0.9222 | 0.8806 | 0.989 | 0.7568 | 0.8956 | 0.9311 | 0.9798 |
|  | Weighted median | 1.0448 | 1.0773 | 0.9877 | 0.9986 | 0.9716 | 0.9777 | 0.9765 |
|  | MRMix | 0.712 | 0.719 | 0.717 | 0.7195 | 0.889 | 0.801* | 0.81228 |
| IL2ra | MR-RAPS<br>(shrinkage) | 0.9784 | 0.9899 | 0.9887 | 0.9452 | 1.0141 | 1.0487* | 1.059** |
|  | MR-Egger | 0.8971 | 0.8021 | 0.8679 | 0.9056 | 0.9555 | 0.9296 | 0.8668 |
|  | Weighted median | 1.0496 | 1.0673 | 1.0425 | 1.0116 | 1 | 1.0079 | 0.9871 |
|  | MRmix | 1.248 | 1.235 | 1.301 | 0.939 | 1.365 | 1.4141 | 1.414 |
| IL18 | MR-RAPS<br>(shrinkage) | 0.925 | 0.918 | 0.95 | 0.925 | 0.952 | 0.972 | 0.974 |
|  | MR-Egger | 0.5528 | 0.597 | 0.659 | 0.697 | 0.79 | 1.006 | 1.023 |
|  | Weighted median | 0.923 | 0.922 | 0.939 | 0.933 | 0.952 | 0.959 | 0.966 |
|  | MRMix | 1.292**** | 1.021** | 0.952 | 0.806 | 1.239 | 1.1727 | 1.31 |

\*:P < 0.05; \*\*: P < 0.001; \*\*\*:P < 0.00001 \*\*\*\*: P < 0.000001

Of nine cytokines, IL18 showed the strongest positive causal link with schizophrenia at the IV selection threshold of 5 × 10^−8^ (OR = 1.292, P = 8.37 × 10^−42^), followed by TNFa at the IV threshold of 1 × 10^−6^ (OR = 0.721, P = 7.33 × 10^−6^), indicating the possible core roles played by these two cytokines in the immune response induced by schizophrenia. The majority of SCZ-induced cytokines generated nominal significance using both MR-RAPS and MRMix, except IL18, IL2ra, and IP10. Of note, the causal estimates of all cytokines were weak-to-modest in effect size with a certain degree of inconsistency existing within the estimate of cytokine at different IV threshold, possibly arising from the sample heterogeneity.

### Two cytokine clusters emerging from the SCZ-induced cytokines

The genomic SEM of commonality underlying nine SCZ-induced cytokines failed to converge, indicative of an ill-fitting model. The subsequent agglomerative hierarchy clustering analysis grouped nine cytokines into three clusters (Figure 3A), the module assignment survived the bootstrapping of 10000 times (Figure 3B, 3C). IL9 and IL2ra were clustered together as module A while SCGFb, IP10, CTACK and IL6 were grouped together as module B, with IL18, GROa and TNFa clustering together as module C. Using the composite summary statistics for each module as the source of IVs, the second round of MR indicated that module A (IL9-IL2ra) did not show any causal relationship with IDPs which could survive the stringent Bonferroni correction (0.05/627 = 7.974e-5); whereas module B (SCGFb-IP10-CTACK-IL6) was found to significantly increase the level of IDPs including IDP_T1_SIENAX_peripheral_grey_normalised_volume (β = 0.0453, P = 4.40×10^−10^, Figure 4A), IDP_dMRI_TBSS_MD_Posterior_corona_radiata_R (β= 0.0584, P = 8.89×10^−16^, Figure 4B) and IDP_dMRI_TBSS_MD_Cingulum_hippocampus_R (β = 0.0563, P = 9.81×10^−15^, Figure 4C); furthermore, module C (IL18-GROa-TNFa) could significantly increased the level of IDP_dMRI_TBSS_L2_Posterior_thalamic_radiation_R (β= 0.0341, P = 2.67×10^−6^, Figure 4D). Intriguingly, no significant direct causal link could be built between the schizophrenia under the different strength level of IVs (5 × 10^−8^, 1 × 10^−5^, 0.001) and these identified IDPs, hinting at a critical mediating role played by the cytokine modules in the pathophysiology of schizophrenia.

**Figure 3.**
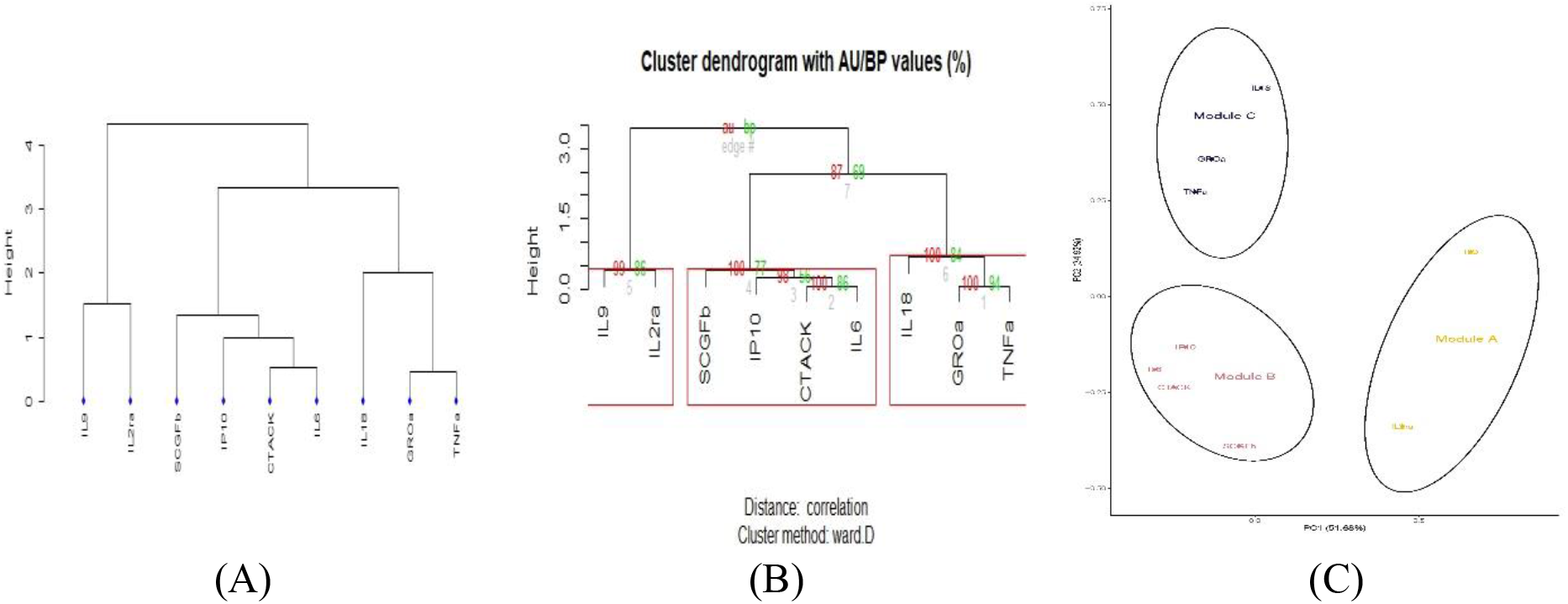
Clustering assignment of nine SCZ-induced cytokines

**Figure 4.**
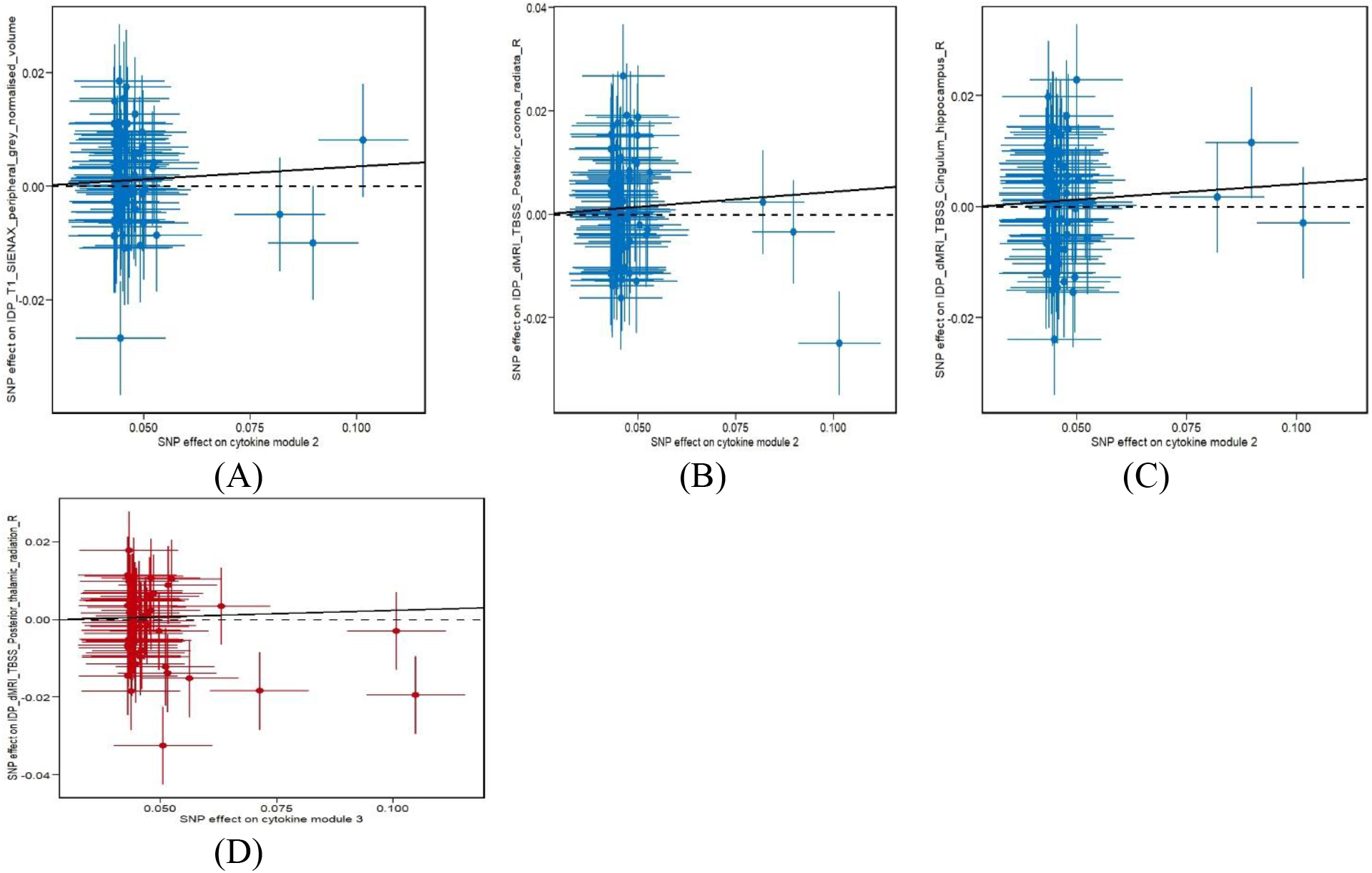
(A-C) Genetic associations with risk factor (cytokine module 2) and outcome (from left to right: IDP_T1_SIENAX_peripheral_grey_normalised_volume, IDP_dMRI_TBSS_MD_Posterior_corona_radiata_R, IDP_dMRI_TBSS_MD_Cingulum_hippocampus_R) for 78 genetic variants as IVs; the blue lines are 95% confidence intervals for the genetic associations (all associations are oriented to the module 2– increasing allele); (D) Genetic associations with risk factor (cytokine module 3) and outcome (IDP_dMRI_TBSS_L2_Posterior_thalamic_radiation_R) for 61 genetic variants as IVs; the red lines are 95% confidence intervals for the genetic associations (all associations are oriented to the module 3-increasing allele);

## Discussion

Our current study tried to detangle the immune responses, proxy by cytokines, of schizophrenia, and the effect of the responses herein on the brain structure and function. The results showed that nine cytokines (SCZ-induced cytokines) were potentially involved in the downstream immune cascade of schizophrenia. Further, three modules could be established from nine SCZ-induced cytokines. Subsequent scanning of the possible roles the three modules in brain structure and function pointed to the causal association between module B and module C to white matter fiber mainly in the mesolimbic system and peripheral cortical gray matter volume.

From the perspective of methodology, we chose a two-sample MR method to delve into the immune component of schizophrenia. The two-sample MR has been widely applied to the establishment of causality between phenotypes. Instead of only choosing SNPs of genome-wide significance as IVs, which explaine a limited proportion of phenotypic variance, thus underpowered in the study of complex traits. We adopted a similar approach as that of PRS (polygenic risk score) analysis, *i.e., calculating the* cumulative causal effect of schizophrenia on cytokines, which is consistent with the polygenic background of schizophrenia. We argue that the approach could capture the immune alterations in schizophrenia in the broadest way. Indeed, in our first round of MR analysis (schizophrenia ⇒ cytokines), the trend of cumulative causality differs notably between cytokines, the study by Brodin et al. assessed the heritability of immune system including cytokines in a Swedish twin population and demonstrated that the serum cytokines varied considerably in the MZ concordance rate. While the heritability of cytokine is beyond the scope of our current study, our present study echoed the findings from Brodin et al. that while the heritability plays a role in how immune system responds to the exposure risk, a heterogeneity exists in the response from different cytokines, which underscores the adaptive nature of immune system. Besides, a certain degree of variability was detected within cytokines using different methods at different selection threshold of IVs. The fact that causal estimates range around 1 existed widely in MR analysis, which suggests the different emphasis methods put on various aspects of IV and the possible effect of confounders such as environmental risks(Burgess, Foley, & Zuber, 2018). A future study could focus on the consensus approach incorporating the strength of different methods.

At the same time, effect size of MR analysis was modest for all investigated cytokines, which is consistent with previous studies proposing a low-grade inflammatory network of multiple cytokine/chemokine with small-to-modest contribution underpins the pathophysiology of schizophrenia (Lesh et al., 2018; Zakharyan & Boyajyan, 2014). Of these nine SCZ-induced cytokines, the causal estimate of IL18 generated the most robust significant P value. IL18 belongs to the IL1 superfamily, this proinflammatory cytokine is secreted by various type of cells including those in the brain, such as pituitary gland, ependymal cell, the neurons of the medial habenula and astrocytes in cerebellum (Alboni, Cervia, Sugama, & Conti, 2010; Tsutsumi et al., 2014). Different lines of evidence suggested the possibility that IL-18 may be one mediator of behavior (Alboni et al., 2009; Zorrilla et al., 2007), cognition(Yaguchi, Nagata, Yang, & Nishizaki, 2010) and neuropsychiatric conditions (Younger, Chen, Hong, Chen, & Tsai, 2003). One major issue regarding the interpretation of these previous studies is the difficulty in determining the action direction of IL18, i.e. whether the change in IL18 level contributes to these pathologies or it is a consequence of the disorders. Our study, for the first time, provided evidence of action direction from schizophrenia to IL18. Further, IL18 was clustered with TNFa and GROa/CXCL1 in module 3 in our analysis. Both TNFa and IL18 was found to induce neurotoxicity through elevated glutamate production which lead to neuronal death (Ye et al., 2013), possibly mediating by GROa.

The signal of IP10/CXCL10 (denoted as IP10 hereafter), an interferon (IFN)-λ inducible protein, is the strongest in width (P-value thresholds). IP10/CXCL10 were found to be involved in many biological functions such as the promotion of the chemotactic activity of CXCR3+ cells, inducing apoptosis, regulating cell growth (Liu et al., 2011). In the studies comparing the serum levels of IP10 between patients with schizophrenia and healthy controls, the results were inconsistent (Asevedo et al., 2013; Hong et al., 2017), which might be due to sample size or heterogeneity in the disease group. Our study, for the first time and based on the data from a large sample, identified the directed effect of schizophrenia on IP10. In the clustering analysis, IP10 was enriched in module B with SCGFb, CCL27, and IL6. One study showed that both CCL27 and IL6 responded to chronical psychosocial stress (Polacchini et al., 2018), and these four cytokines were enriched in the brain hypothalamus (https://fuma.ctglab.nl/gene2func) with a nominal significant P value. Our study provided another piece of evidence regarding the synergized role of module B in the pathology of schizophrenia.

Previous neuroimaging studies have generated inconsistent conclusions about the deviation of cortical gray matter volume and white matter fibers in patients with schizophrenia (Haijma et al., 2012). The reason for such an inconsistency could be the modest sample size and inherent heterogeneity of the disorder. Although future study to assess the expression level of these cytokine genes in the brain tissue of schizophrenia is required, our current findings based on the output of large population-based GWAS indicated that the relationship between these brain structural changes and schizophrenia might be conditioning on some other factors such as personalized inflammatory profiles.

Due to the sample size of cytokine GWAS, we did not carry out the bi-directional MR analysis in the present study. We are fully aware that a complete casual chain is needed to map the immune pathophysiology of schizophrenia, which emphasized a genome-wide study of immune parameters in a larger population in order to capture the more accurate effect sizes of SNPs as IVs.

## Conclusion

In summary, using an approach of two-sample Mendelian randomization and rich resource of GWAS, the current study identified nine cytokines, the highlight of which includes IL18 and TNFa, to be causally associated with schizophrenia. These SCZ-induced cytokines could be clustered into three modules, and the modules represented by IL18 and IP10 could infer an effect on the total gray matter volume and the white matter fibers in the mesolimbic system. Our study, for the first time, mapped the causal link from schizophrenia to the comprehensive immune responses and the findings provided some evidence worth further explorations to further understand the detailed etiological mechanism of schizophrenia.

